# The therapeutic effect of diet and dietary ingredients on cellular senescence in animals and humans: A systematic review

**DOI:** 10.1101/2023.07.28.550928

**Authors:** Lihuan Guan, Anna Eisenmenger, Karen C. Crasta, Elena Sandalova, Andrea B. Maier

## Abstract

**Background:** Cellular senescence is a permanent state of cell cycle arrest and has been regarded as a therapeutic target for ageing and age-related diseases. Several senotherapeutic agents have been proposed, including compounds derived from natural products which hold the translational potential to promote healthy ageing. It is largely unclear whether cellular senescence could be targeted by dietary interventions. This systematic review examined diets and dietary ingredients and their association with cellular senescence load in animal models and humans, with an intent to identify dietary interventions with senotherapeutic potential.

**Methods:** The databases PubMed and Embase were systematically searched for key terms related to cellular senescence, senescence markers, diets, nutrients and bioactive compounds. Intervention and observational studies on human and animal models investigating the effects of diets or dietary ingredients via oral administration on cellular senescence load were included. The studies were screened using the Covidence systematic review software. Study design, methods and results were extracted. Biomaterials used for senescence detection were categorized into physiological systems. The SYRCLE’s risk of bias tool and Cochrane risk of bias tool v2.0 were used to assess the risk of bias for animal and human studies respectively.

**Results:** Out of 5707 identified articles, 82 articles consisting of 78 animal studies and 4 human studies aimed to reduce cellular senescence load using dietary interventions. In animal studies, the most-frequently used senescence model was normal ageing (26 studies), followed by D-galactose-induced models (17 studies). Resveratrol (8 studies), vitamin E (4 studies) and soy protein isolate (3 studies) showed positive effects on reducing the level of senescence markers such as p53, p21, p16 and senescence-associated ß-galactosidase in various tissues of physiological systems. In three out of four human studies, ginsenoside Rg1 had no positive effect on reducing senescence in muscle tissues after exercise. The risk of bias for both animal and human studies was largely unclear.

**Conclusion:** Resveratrol, vitamin E and soy protein isolate are promising senotherapeutics studied in animal models. Studies testing dietary interventions with senotherapeutic potential in humans are limited and translation is highly warranted.

## 1. Introduction

Ageing is characterized by a progressive deterioration in physiological function ubiquitous to all living organisms (Li *et al*, 2021b). Accumulation of cellular damage and loss of function with age affects organismal health, making ageing a primary risk factor for metabolic syndromes, musculoskeletal disorders, cardiovascular diseases and neurodegenerative diseases among others (Salvatore, 2020). Cellular senescence, one of the key ageing hallmarks, is featured as a stable cessation of cell proliferation, macromolecular damage, dysregulated metabolism, and acquisition of a pro-inflammatory and proteolytic secretome, namely senescence-associated secretory phenotype (SASP) (Gorgoulis *et al*, 2019; Lopez-Otin *et al*, 2013). The utility of multiple senescence-associated markers, rather than a sole marker, has been recommended to identify and classify a cell state as senescence (Cohn *et al*, 2023).

Senescent cells accumulate with age (Tuttle *et al*, 2020) and provoke tissue dysfunction, which has been associated with a wide spectrum of age-related diseases (Munoz-Espin & Serrano, 2014; Tuttle *et al*, 2021); thus, researchers focus on senotherapeutics, including senolytics that selectively eliminate senescent cells (i.e. senolysis), and senomorphics, that suppress SASP to promote healthy ageing and prevent or improve age-related conditions (Raffaele & Vinciguerra, 2022). Daily diet interventions from natural products exerted benefits in prevention and treatment for age-related conditions (Dhanjal *et al*, 2020). Compounds derived from natural products have shown to be capable of reducing senescence load and eliciting anti-inflammatory effects *in vitro* and *in vivo* of animal models and humans, demonstrating the potential for utility as senotherapeutics (Deledda *et al*, 2022; Li *et al*, 2019; Zhang *et al*, 2023).

This systematic review aimed to assess the association between dietary intake and cellular senescence load in animal models and human subjects for the identification of diets and/or dietary ingredients with senotherapeutic potential that could prove beneficial for optimal human health during ageing.

## 2. Methods

### 2.1 Information sources and search strategy

The systematic review was registered at PROSPERO International prospective register of systematic reviews (Reg #: CRD42022338885). The databases PubMed and Embase were systematically searched by two reviewers from inception until the 18^th^ of June 2022. The search strategy included keywords, MeSH (Medical Subject Headings) and Emtree (EMBASE subject heading) terms for the respective databases for diets, dietary ingredients, cellular senescence and senescence markers. Database specific filters were used to apply exclusion criteria (**Appendix A: Search strategy**).

### 2.2 Eligibility Criteria

Studies were included according to the following criteria: 1) used interventions including diets and dietary ingredients; 2) measured cellular senescence markers; 3) observational and interventional studies; 4) population: animals and humans aged older than 18 years. The following exclusion criteria were applied: 1) interventions with exclusively Food and Drug Administration controlled drugs and caloric restriction or fasting; 2) dietary interventions not administered through the gastrointestinal tract; 3) conference abstracts, reviews, editorials, letters, books, book chapters and short surveys; 4) articles presenting only methods or theories; 5) corrected articles; 6) human populations with less than five participants; 7) studies not published in English (**Appendix A: Table S1**). *In vitro* studies were excluded, whereas *ex vivo* studies that examined the impact on tissues after the intervention in cell cultures were included. The current review focused on studies aiming to reduce senescence load.

### 2.3. Article selection and data extraction

After removing duplicates, the Covidence systematic review software (Veritas Health Innovation, Melbourne, Australia; available at www.covidence.org.) was used by two independent reviewers (L.G. and A.E.), who first selected the articles by title and abstract screening, followed by the full text screening. Thereafter, data was extracted from included studies. Additional studies were identified by reference screening of included studies. Discrepancies in the study selection and data extraction process were resolved by a third reviewer (K.C or E.S). The variables were independently extracted: first author, year of publication, health condition, age, sex distribution, weight, sample size, intervention, dosage, administration/route, frequency, starting time of intervention, duration, biomaterial, cell type, techniques, markers used to detect senescence (**Appendix B: Table S1**) and outcomes. The statistical significance as either exact or a range of *p*-values was extracted for all outcomes. If reported, mean or median with standard deviation, standard error, confidence intervals, and interquartile range were extracted for continuous outcomes. For binary outcomes, the number of events, total number in groups and percentage of events, or ratios with confidence intervals were extracted. Additional information on species and animal models was extracted from animal studies. Additional information on study design, height, dietary history before and after intervention, washout time and sampling time for outcome measurements was extracted from human studies.

### 2.4 Intervention classification

According to the number and property of ingredients, interventions were classified into diets and diet additives which included multiple ingredients (plant or animal origin and fungus) and single ingredients (nutrients and bioactives) (**Appendix B: Table S2**).

### 2.5 Data analysis

To evaluate the effect of an intervention on senescence, three steps were applied. Step 1: The quality of methodology for each intervention was grouped into three categories according to the number of measured senescence markers and number of studies: a) high, at least two senescence markers used in three or more studies; b) moderate, at least two senescence markers used in one or two studies; c) low, one senescence marker used. Step 2: Each outcome was defined as the change of the given senescence marker measured in the cell type of the given biomaterial by the technique used at each time point, and each dosage. Step3: According to the *p* values and direction of the change in senescence markers in the intervention groups compared to control groups (**Appendix B: Table S1**), each outcome was classified into five types: a) positive, changed in the expected direction (*p* < 0.05); b) positive trend, changed in the expected direction (0.05 < *p* < 0.1 or no *p* values reported); c) insignificant (*p* > 0.1 or no *p* values reported); d) negative trend, changed in the unexpected direction (0.05 < *p* < 0.1 or no *p* values reported); e) negative, changed in the unexpected direction (*p* < 0.05). When multiple SASP markers were measured simultaneously, an overall SASP outcome was given based on the proportion of positive outcomes for each SASP marker: if > 50% SASP markers had positive outcomes, the overall SASP outcome was positive; if 20% - 50% SASP markers had positive outcomes, the overall SASP outcome had a positive trend; if ≤ 20% SASP markers had positive outcomes, the overall SASP outcome was determined by the dominant outcome type. Data were presented as a number X of a specific outcome type out of a total number Y of outcomes (X/Y). Overall, an intervention was considered senotherapeutic if it was investigated with high quality of methodology and if more than half of the outcomes were positive.

The interventions were cross checked with the DrugAge database, a database of drugs, compounds and supplements (including natural products and nutraceuticals) with properties that extend longevity in model organisms (Barardo *et al*, 2017) (https://genomics.senescence.info/drugs/index.php).

### 2.6 Risk of bias

The SYRCLE’s risk of bias tool (Hooijmans *et al*, 2014) and the Cochrane risk-of-bias tool v2.0 (Higgins *et al*, 2011) were used by two independent reviewers (L.G. and A.E.) to assess the bias of animal studies and human studies respectively. Conflicts were resolved by a third reviewer (E.S.). Seven key sources of bias were assessed: sequence generation, allocation concealment, blinding of participants and personnel, blinding of outcome assessment, incomplete outcome data, selective outcome reporting and other sources of bias. Baseline characteristics, random housing and random outcome assessment were additionally assessed for animal studies. The key bias sources were denoted ‘high risk’ or ‘low risk’ depending on if the aspect was deemed to encourage or mitigate bias and ‘unclear risk’ if the aspect was not reported. The overall risk of bias in a study was classified as ‘low’ if all key domains were ‘low risk’. It was classified as ‘high’ or ‘unclear’ if one or more of the domains were either ‘high risk’ or ‘unclear risk’ respectively (Higgins *et al*., 2011).

## 3. Results

### 3.1 Study selection

**Fig 1** demonstrates the study selection process for the association analysis between diet and dietary ingredients, and cellular senescence. The literature search retrieved 9181 articles. Duplicates were removed, leaving 5707 articles for title and abstract screening and 259 articles for full text screening. Out of 121 articles eligible for inclusion, 82 studies described interventions aimed at reducing cellular senescence and 39 studies described interventions aiming to induce cellular senescence.

**Figure 1.**
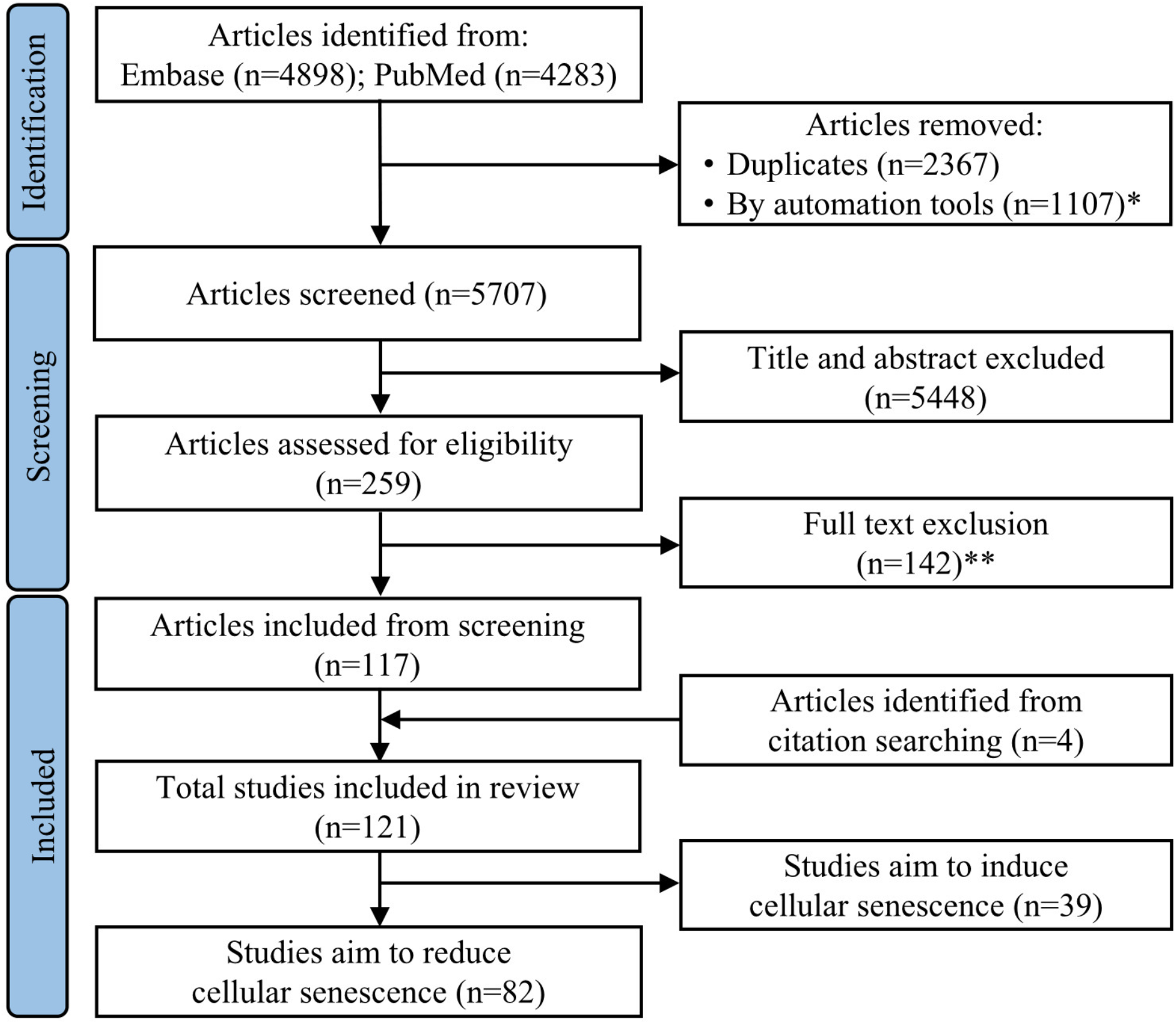
Flow chart of search strategy. *Books, book chapters, conference abstracts, editorials, letters, reviews and short surveys. **Wrong route of administration (n=55), wrong outcomes (n=26), wrong intervention (n=33), wrong study design (n=5), review (n=15), conference abstract (n=7), not in English (n=1).

### 3.2 Study characteristics

Study design and results are provided for animal studies (**Appendix C: Table S1, Table S2**) and human studies (**Appendix C: Table S3, Table S4**), stratified by the type of intervention.

Out of the 82 included studies, 78 studies were conducted in animals and four studies were conducted in humans. Sixty-five out of 78 studies reported that sex and male animals were used most often (52 studies). Animal species included mice (55 studies), rats (22 studies), fish (2 studies) (Liu *et al*, 2018; Xia *et al*, 2014) and hamsters (1 study) (Qi *et al*, 2021). The most-utilised senescence model was normal ageing (26 studies), followed by D-galactose-induced models (17 studies), transgenic models (17 studies), high-fat diet models (9 studies), irradiation (3 studies) (Li *et al*, 2018; Sim *et al*, 2019; Zhang *et al*, 2021a) and ovariectomy (3 studies) (Chen *et al*, 2017b; Zhang *et al*, 2013, 2014). The senescence load in the entire body (Xia *et al*., 2014) as well as in tissues of nine physiological systems consisting of 53 different biomaterials was reported (**Appendix B: Table S3**). Notably, the most investigated biomaterials were the liver (15 studies), aorta (13 studies), hippocampus (10 studies) and brain (8 studies). Most studies reported at least two senescence markers (62 studies). A total of 16 different senescence markers were reported. The cell cycle regulators p53 (44 studies), p16 (44 studies) and p21 (43 studies) and SA-ß-gal (43 studies) were used most frequently as senescence markers (**Table 1**).

**Table 1.**
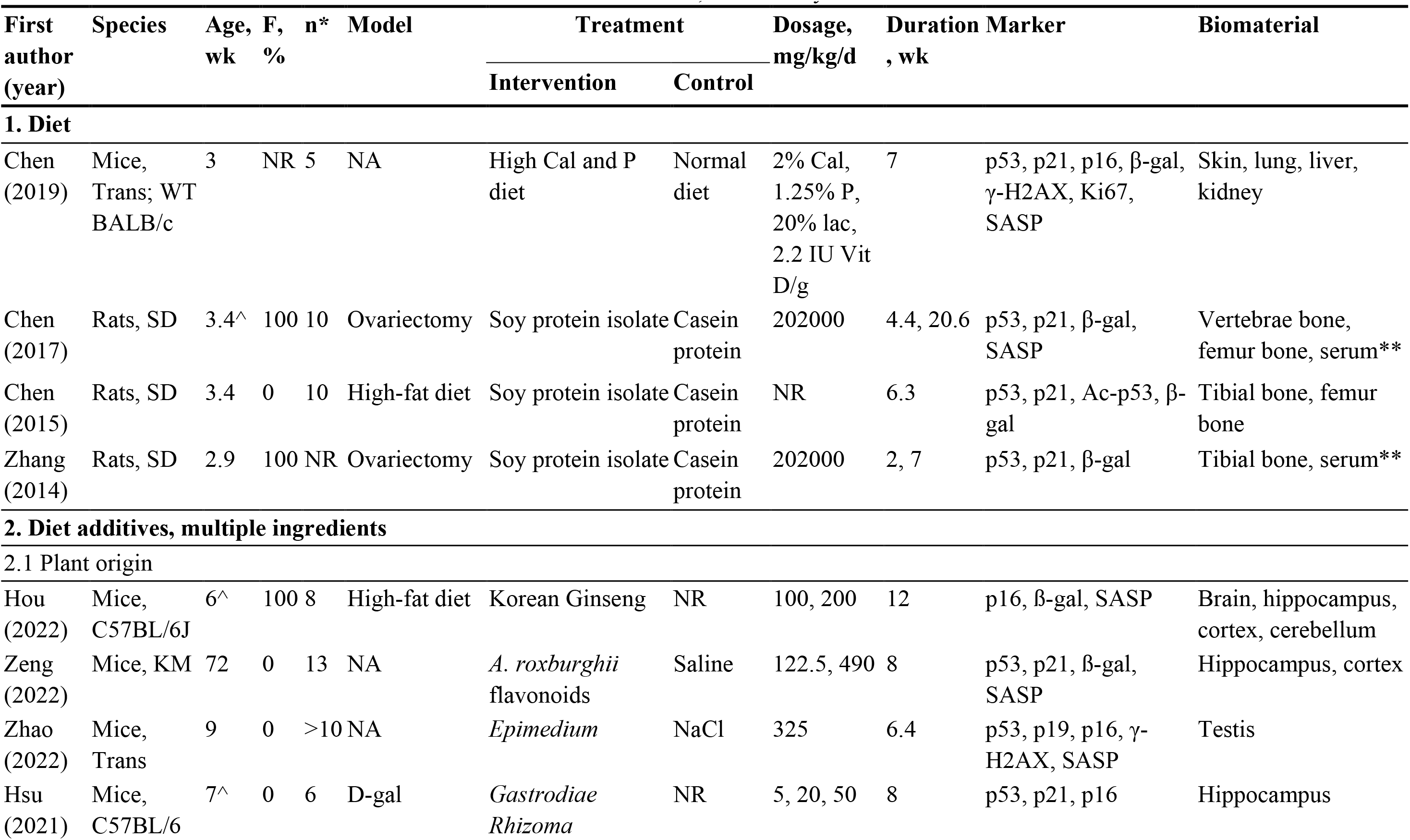
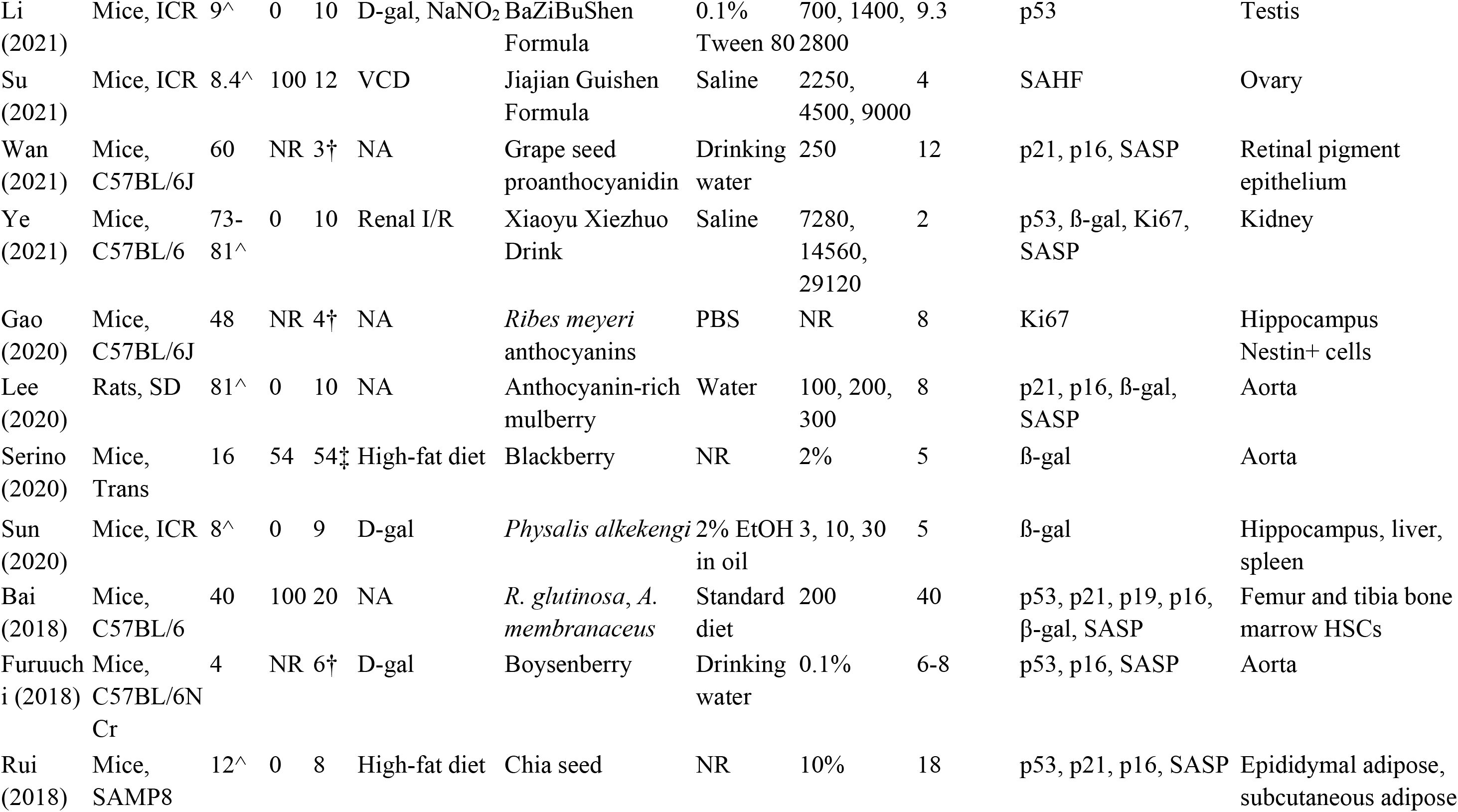
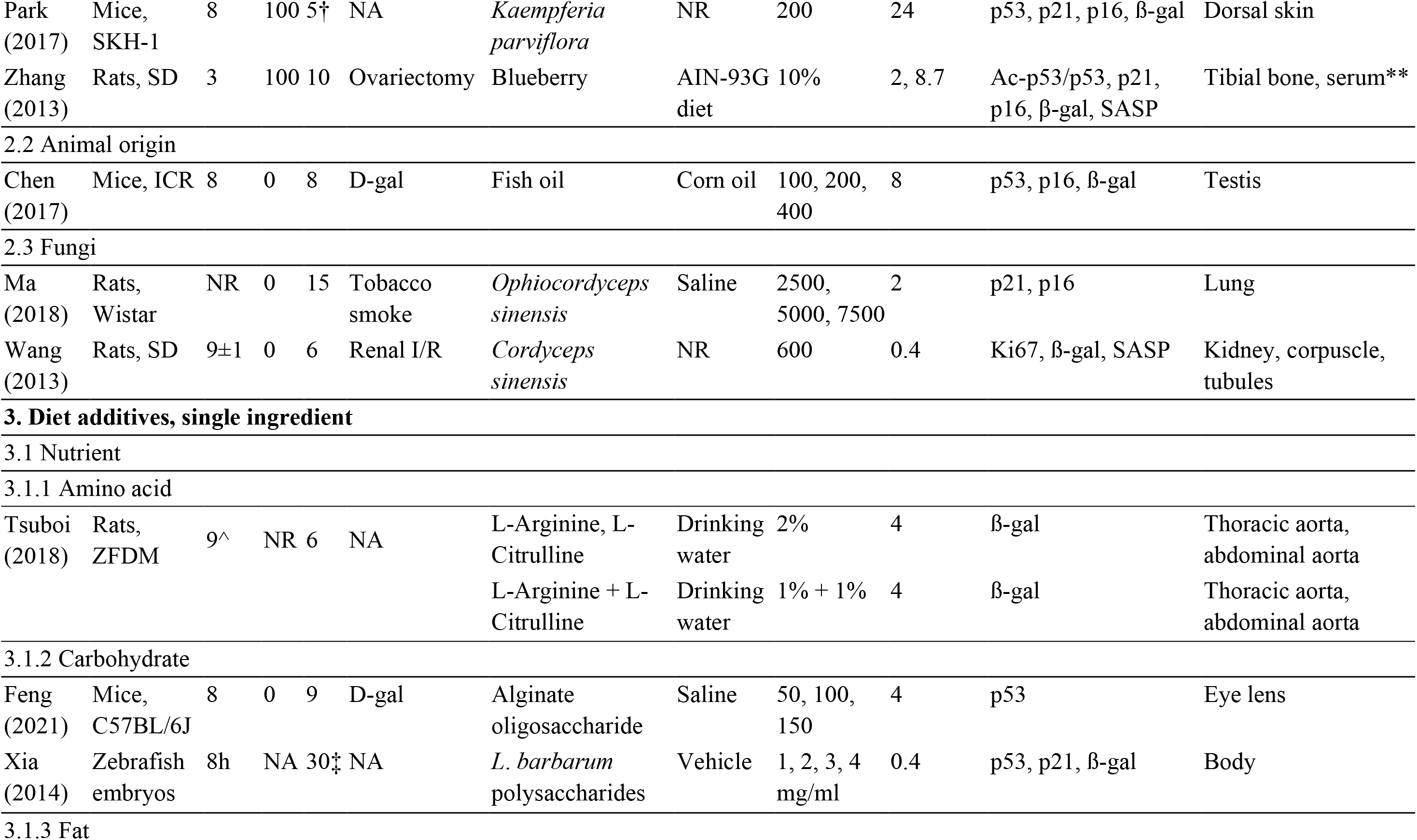
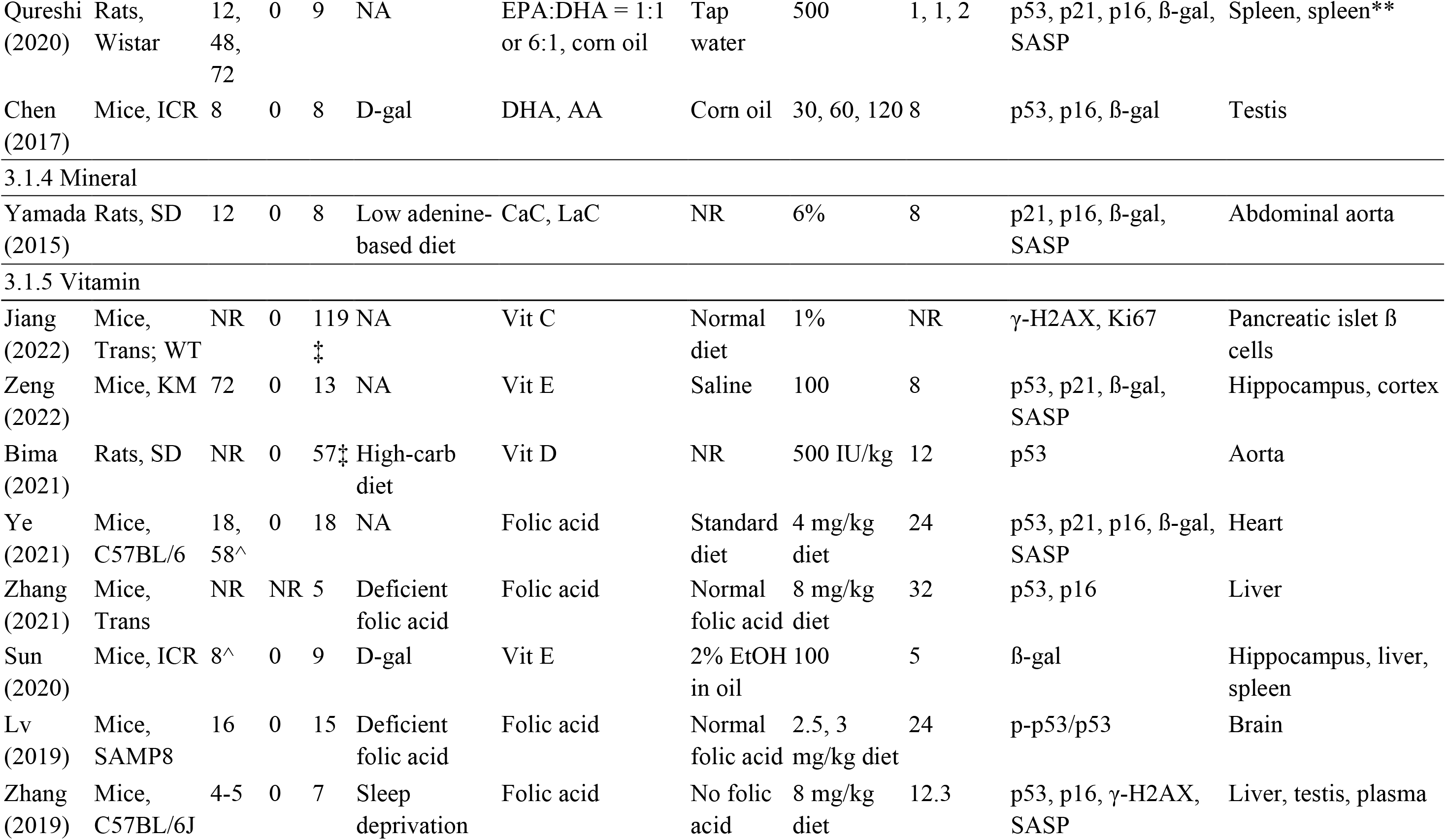
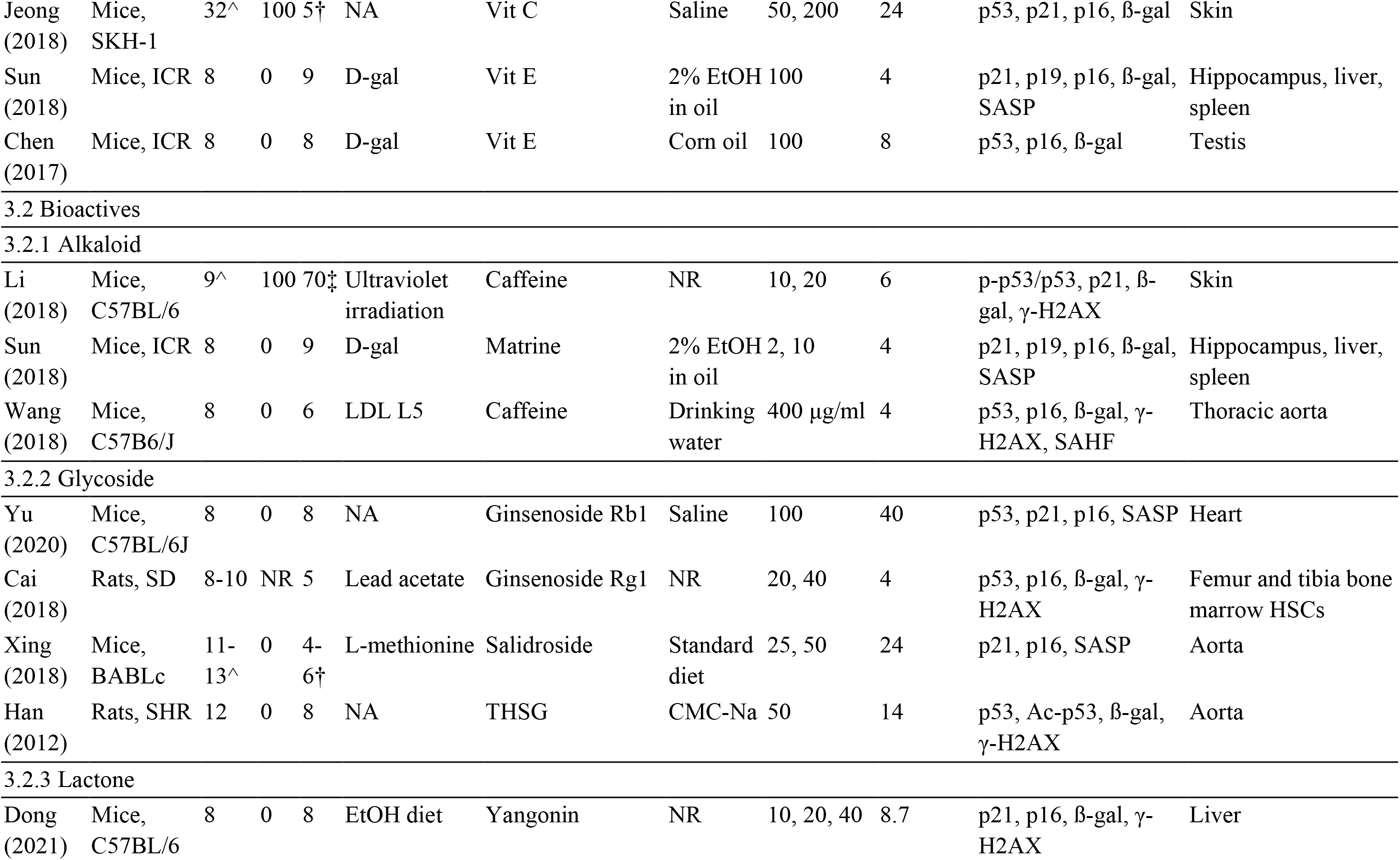
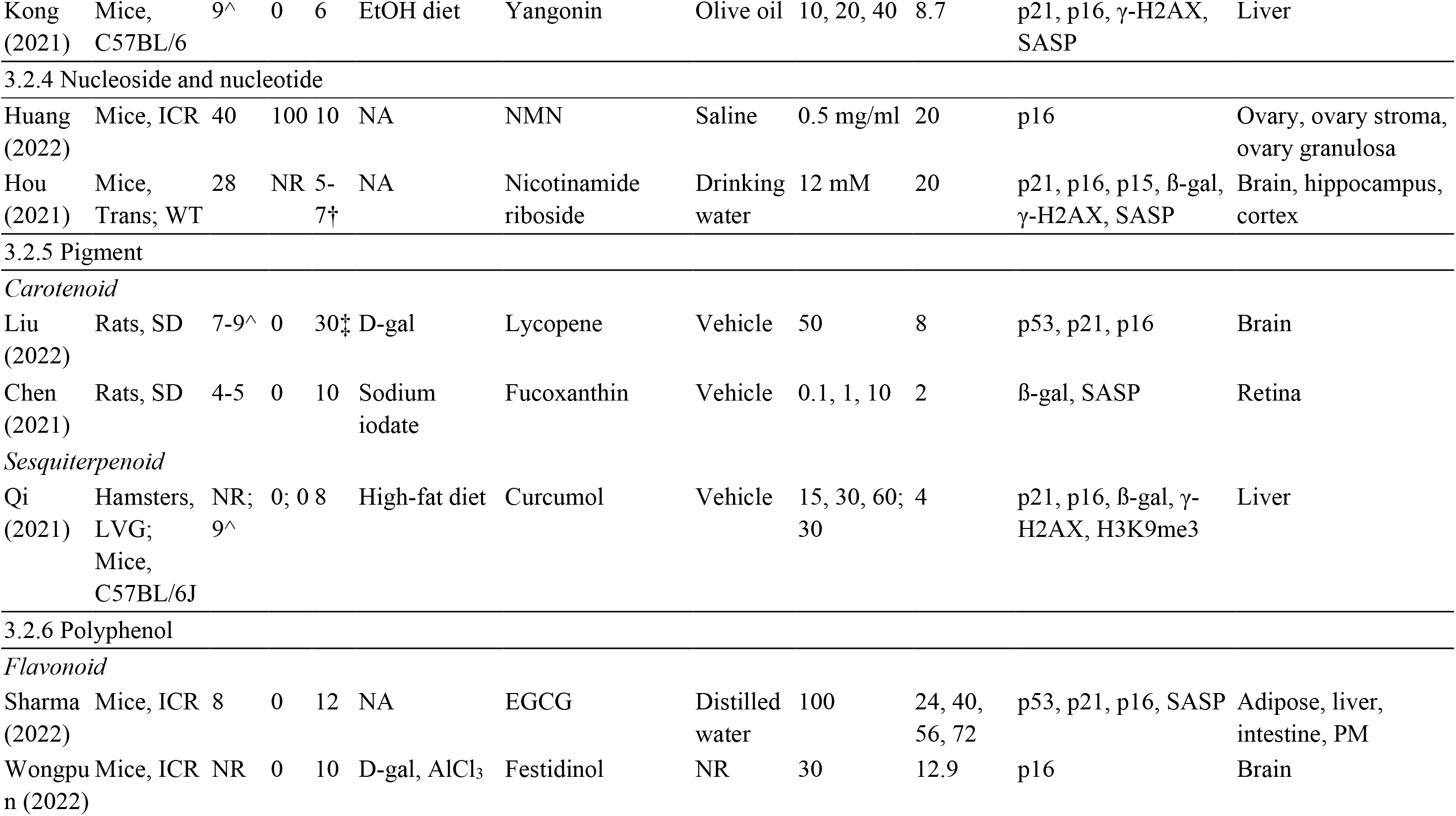
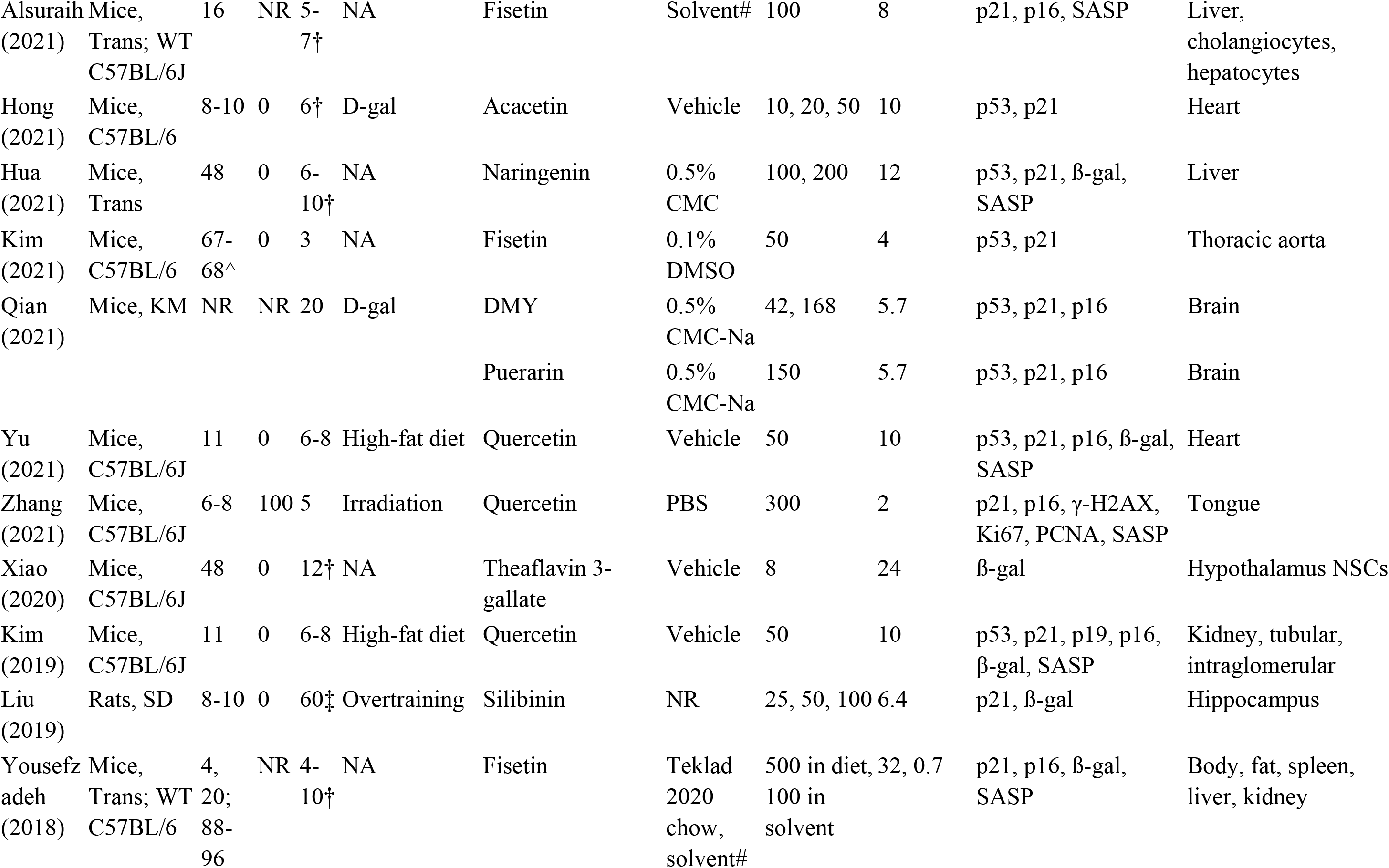
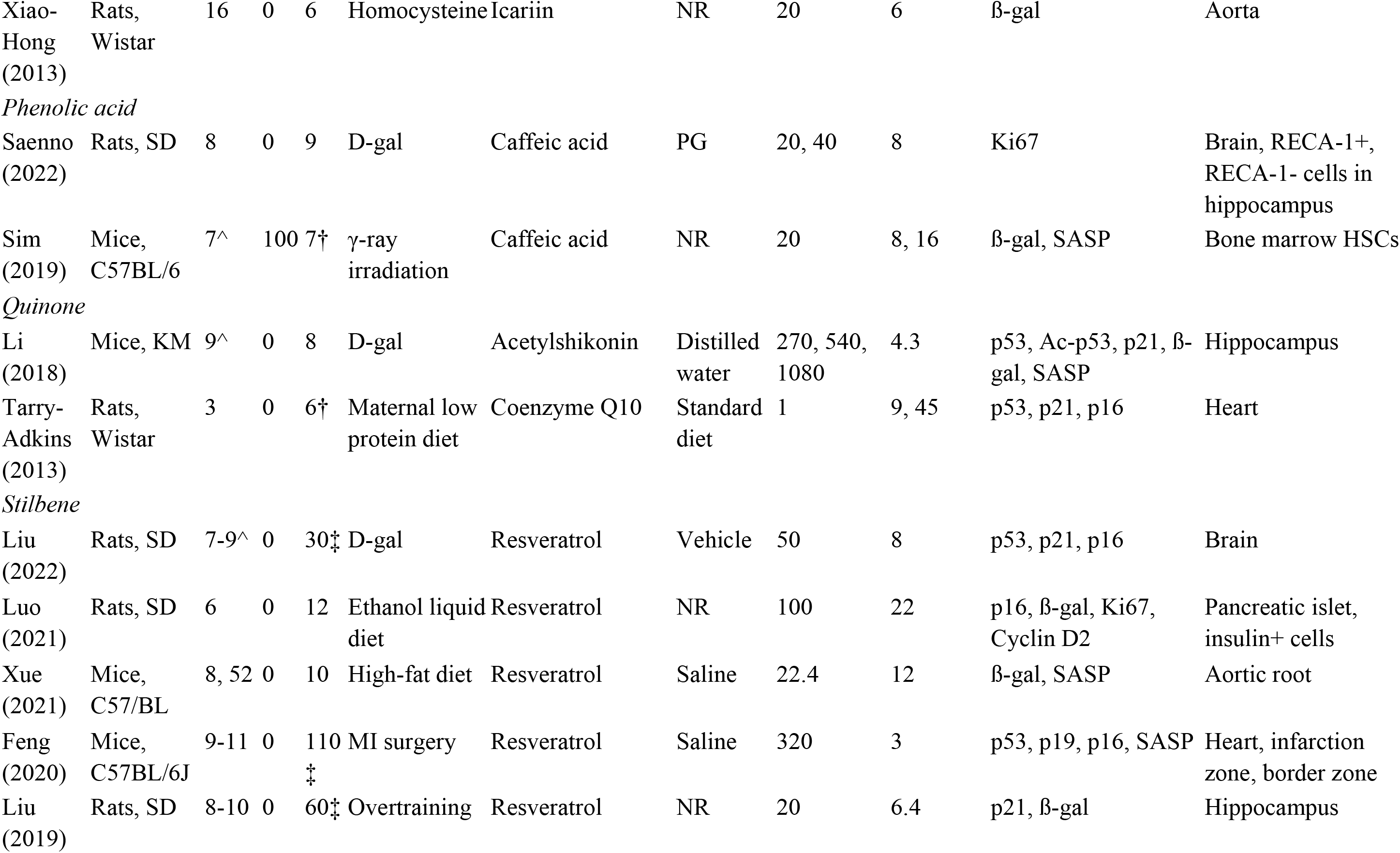
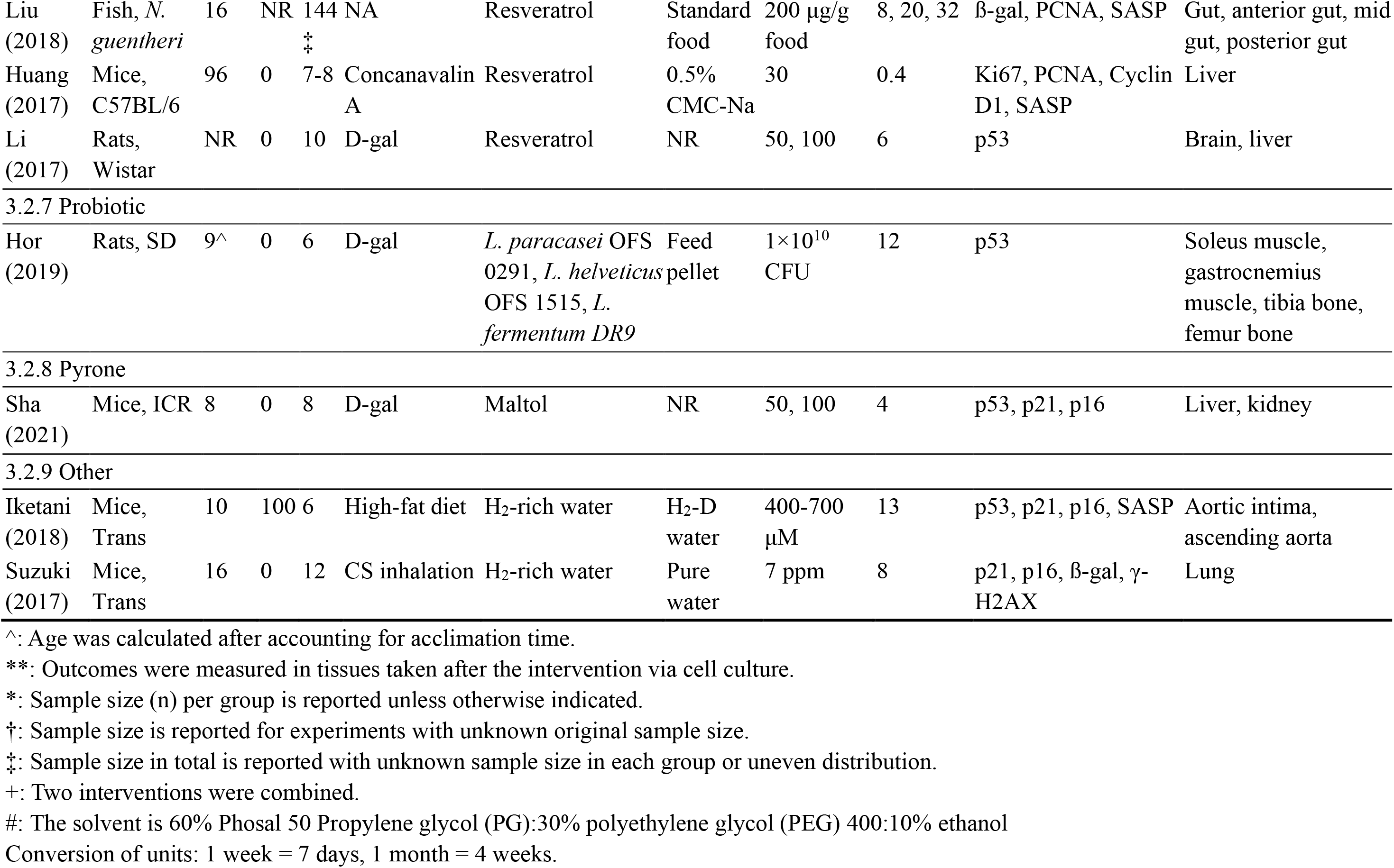
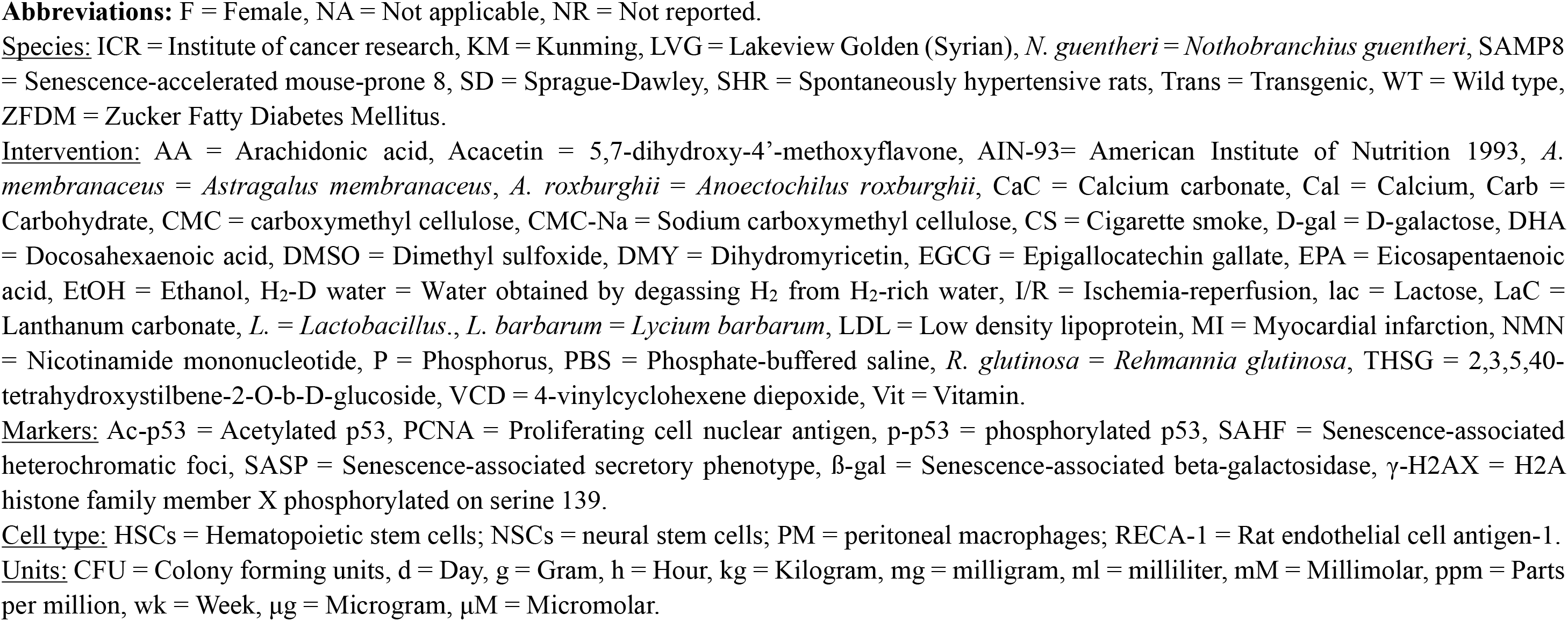
Characteristics and outcomes of intervention studies in animals, stratified by the classification of interventions.

All four human studies are randomized controlled trials conducted in twelve healthy male participants aged 20-25 years old each. The acute bout of resistance exercise that induces muscle damage and hypertrophy was used as a model and senescence load was examined in the vastus lateralis muscle. The cell cycle regulator p16 was used to measure senescence in all four studies (**Table 2**).

**Table 2.**
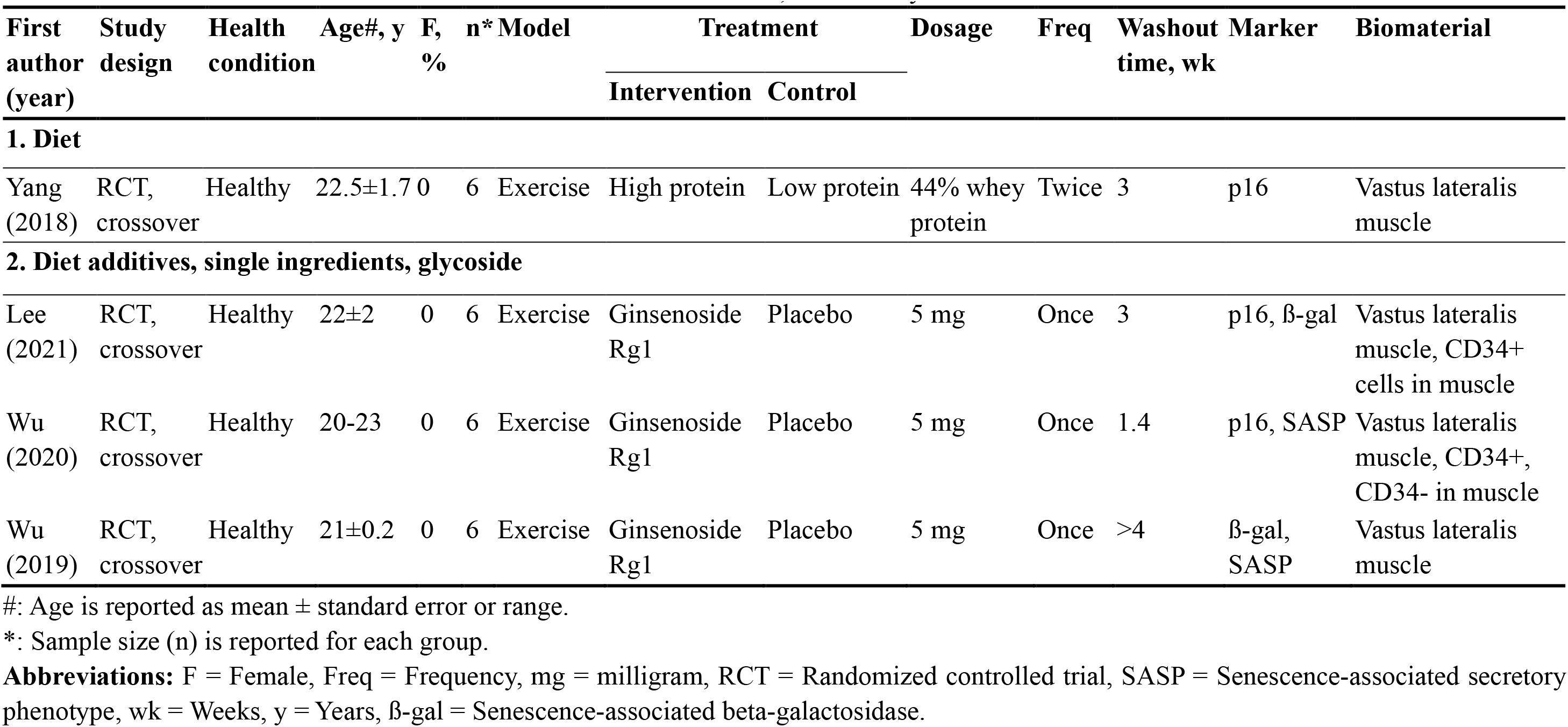
Characteristics and outcomes of intervention studies in humans, stratified by the classification of interventions.

### 3.3 Diet

Soy protein isolate was investigated with high quality of methodology, showing 26 positive outcomes out of 47 (26/47) in ovariectomy models and high-fat diet models (3 studies) (Chen *et al*, 2015; Chen *et al*., 2017b; Zhang *et al*., 2014). The effect of high calcium and phosphate diet on seven senescence markers was investigated across four physiological systems, demonstrating 10/18 positive outcomes in the vitamin D deficiency transgenic model (1 study) (Chen *et al*, 2019) (**Table 3**).

**Table 3.**
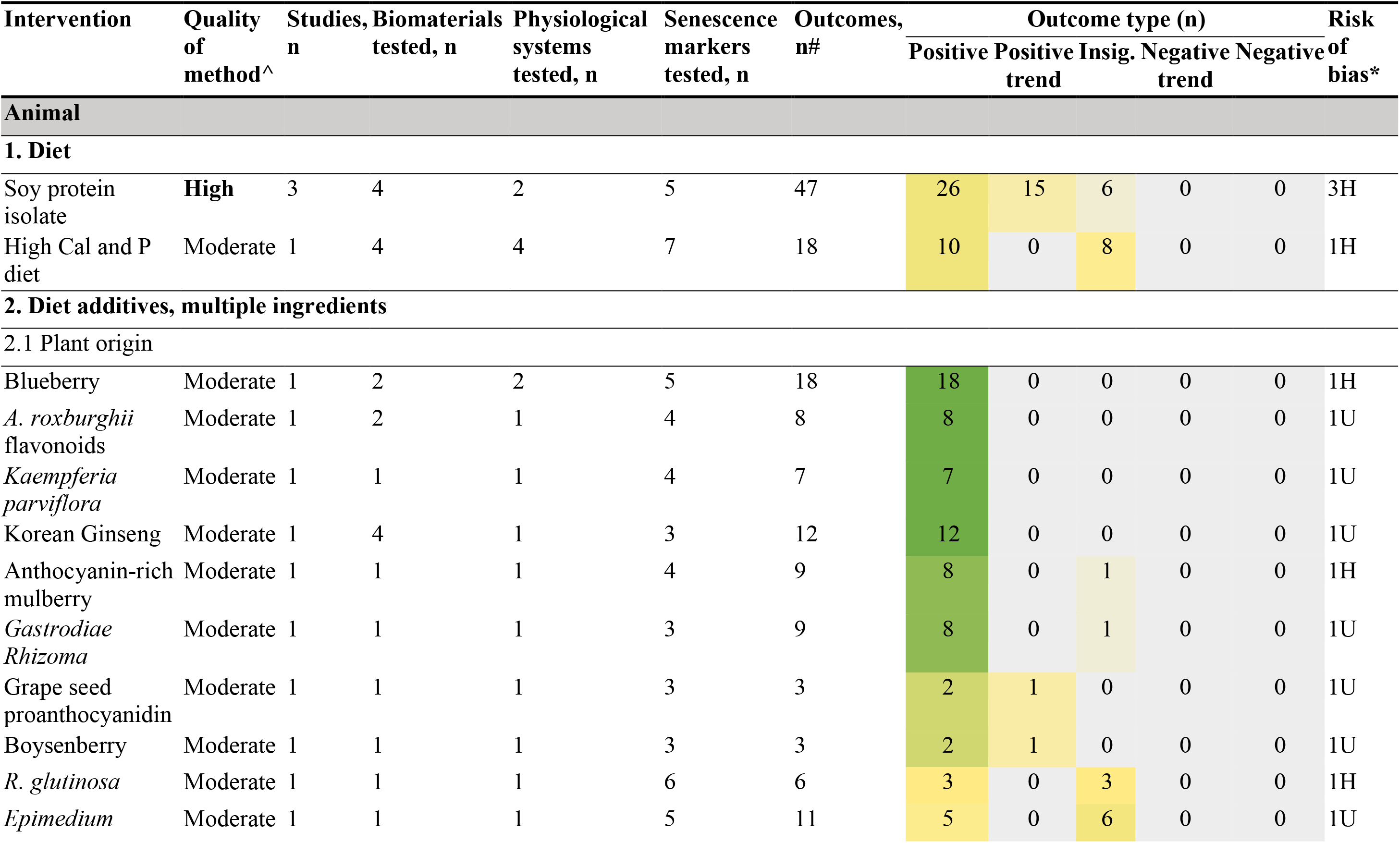
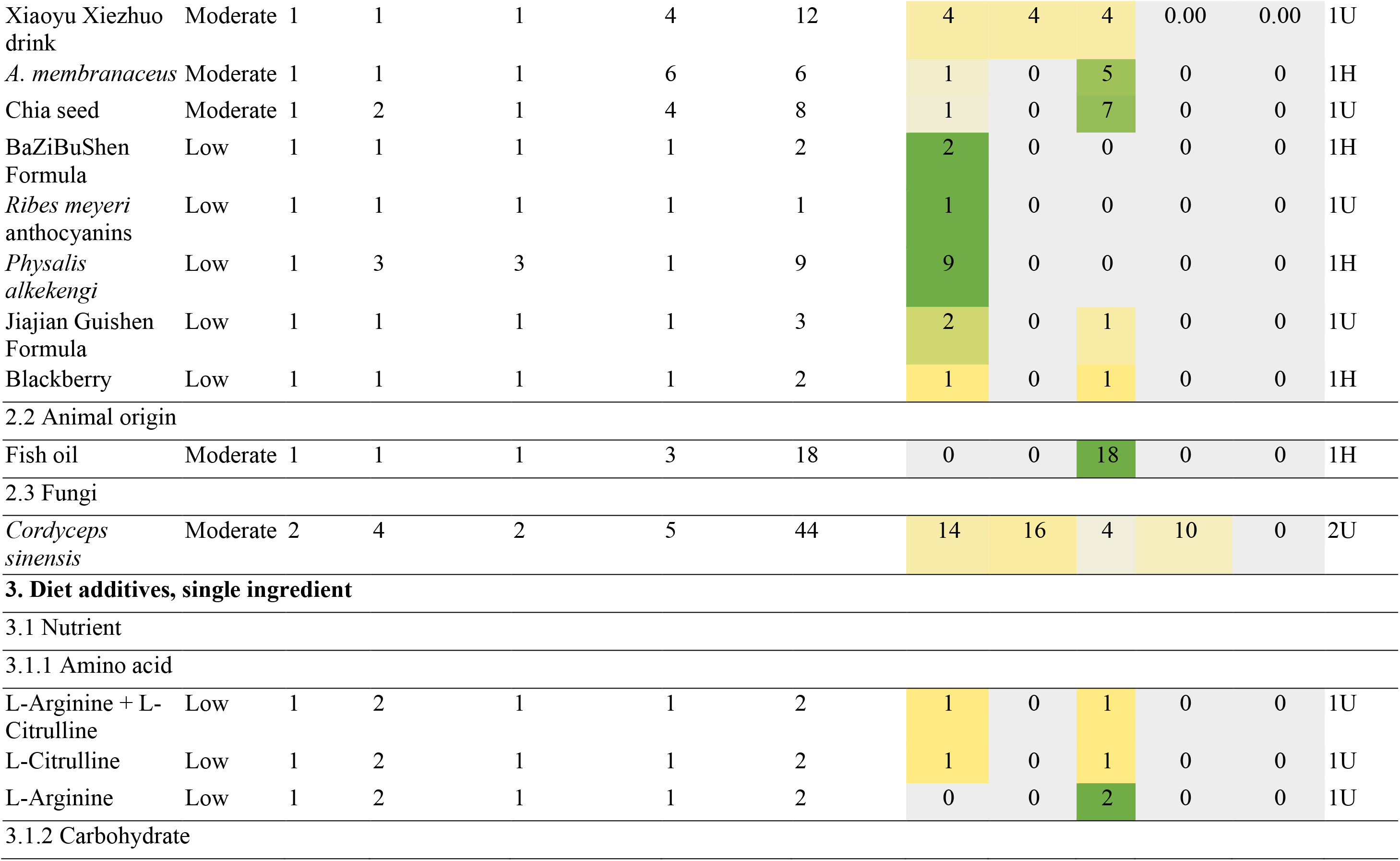
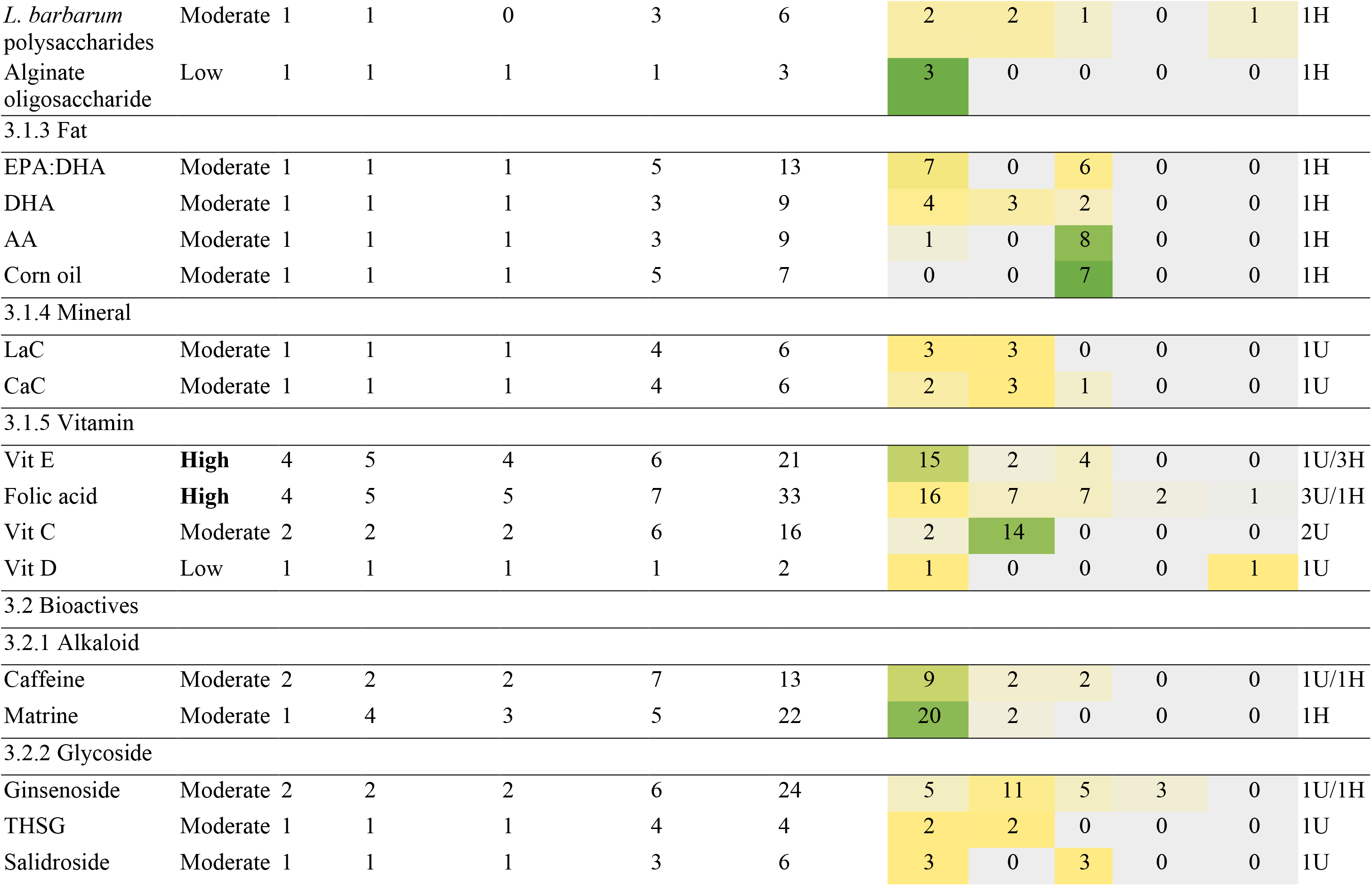
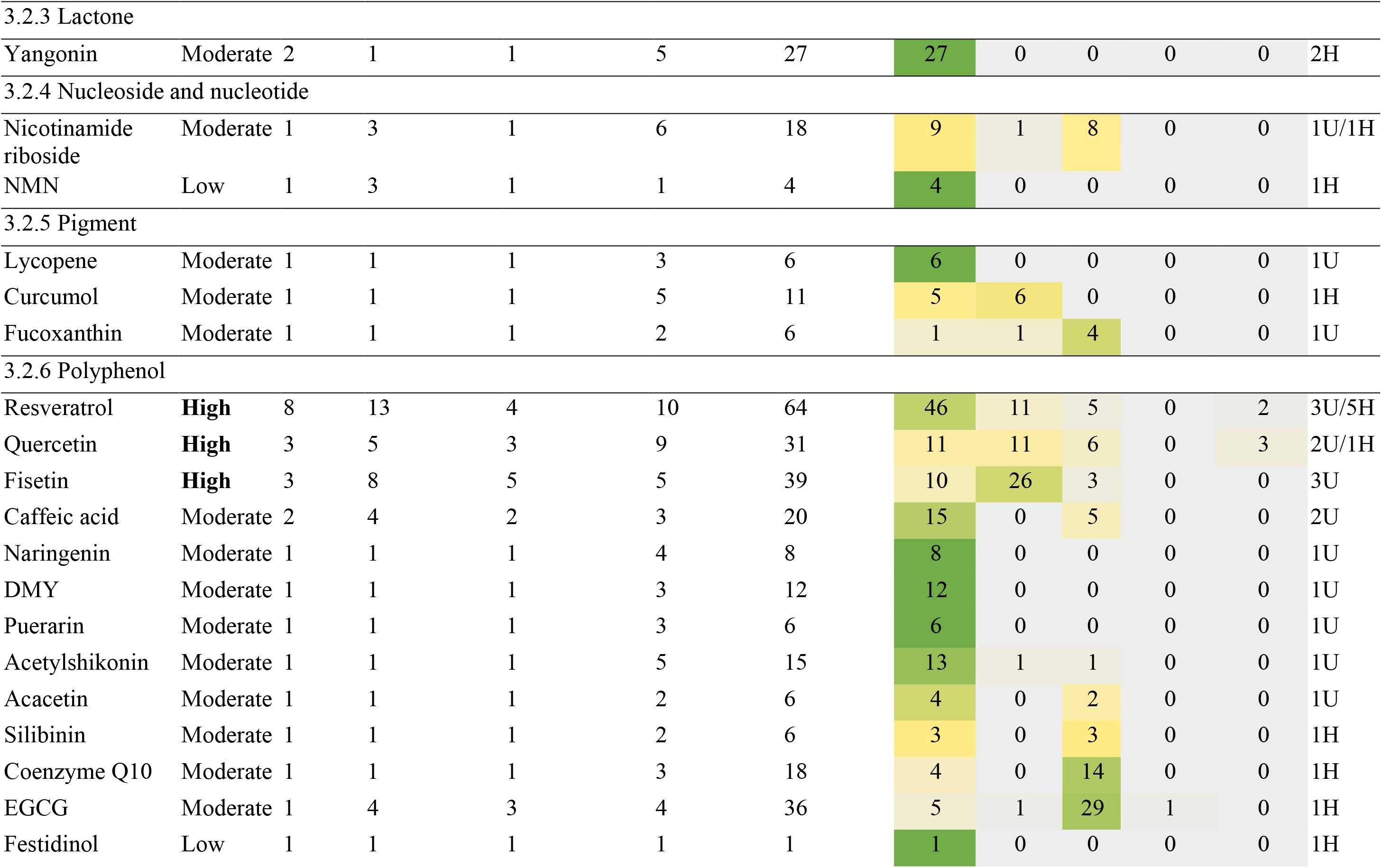
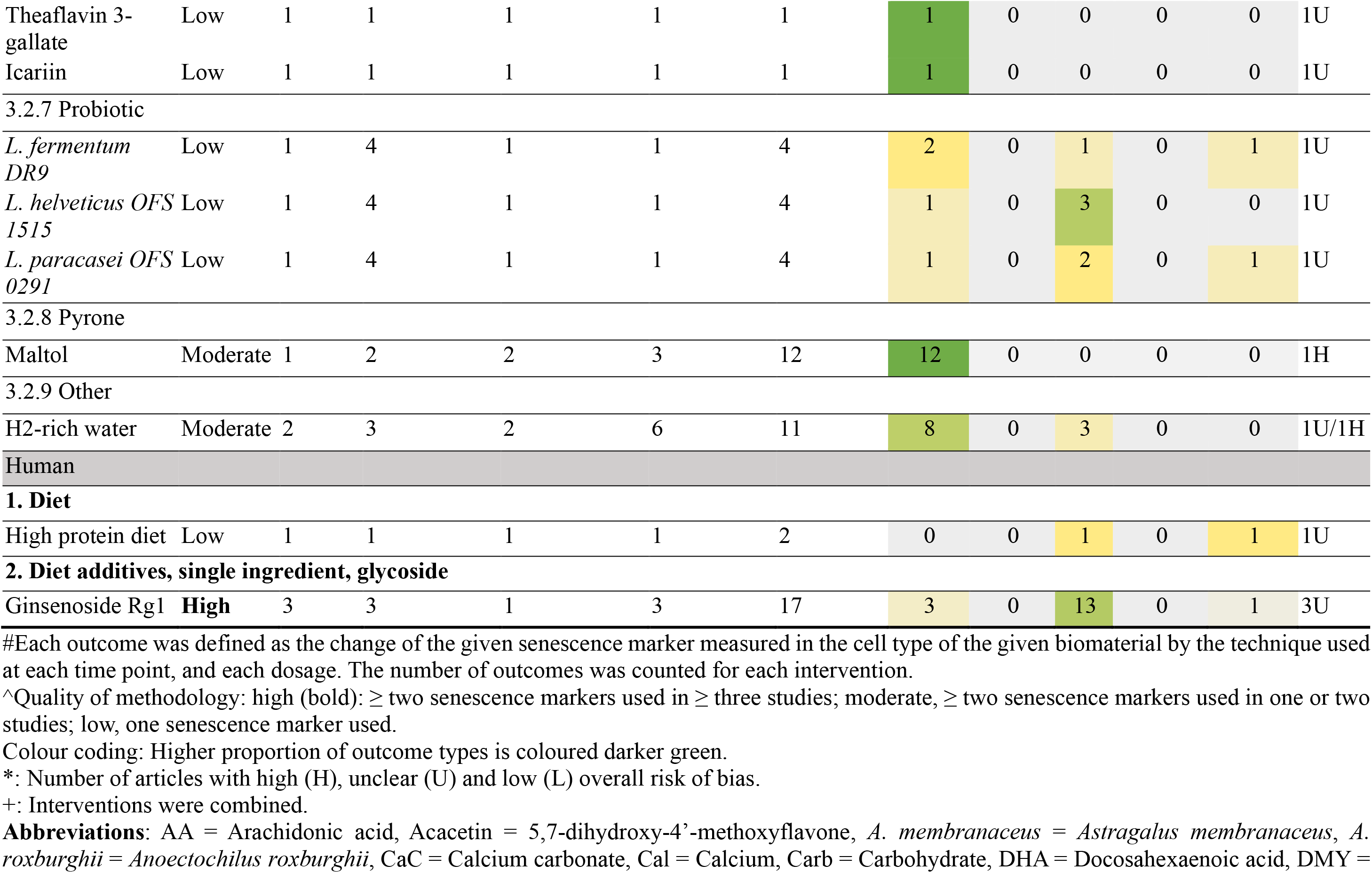
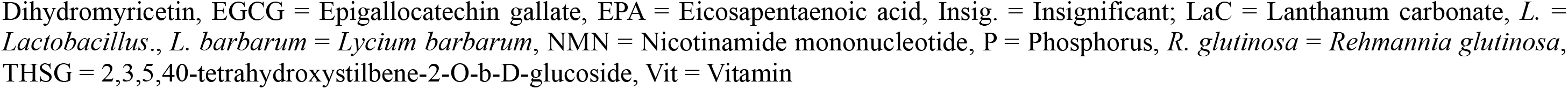
Summary of study characteristics and outcomes of each intervention in animals and humans, stratified by the classification of interventions.

### 3.4 Diet additives, multiple ingredients

#### 3.4.1 Plant origin

Eighteen interventions originated from plants were investigated (17 studies, two interventions were investigated in the same study) (**Table 3**). Normal ageing (6 studies) and D-galactose-induced ageing models (4 studies) were assessed the most. Thirteen interventions had moderate quality of methodology, among which blueberry (18/18) (Zhang *et al*., 2013), *Anoectochilus roxburghii* flavonoids (8/8) (Zeng *et al*, 2022), *Kaempferia parviflora* (7/7) (Park *et al*, 2017), and Korean ginseng (12/12) (Hou *et al*, 2022) showed all positive outcomes. *Epimedium* (6/11) (Zhao *et al*, 2022), *Astragalus membraneceus* (5/6) (Bai *et al*, 2018) and Chia seed (7/8) (Rui *et al*, 2018) showed insignificant outcomes. Studies with low quality of methodology showed that the outcomes from BaZiBuShen formula (2/2) (Li *et al*, 2021a), *Ribes meyeri* anthocyanins (1/1) (Gao *et al*, 2020) and *Physalis alkekengi* (9/9) (Sun *et al*, 2020) were positive.

#### 3.4.2 Animal origin

Fish oil showed no effect on senescence load induced by D-galactose in endocrine systems with 18/18 insignificant outcomes (1 study) (Chen *et al*, 2017a) (**Table 3**).

#### 3.4.3 Fungi

Studies with moderate quality of methodology revealed that *Cordyceps sinensis* had 14/44 positive outcomes with 16/44 outcomes indicating a positive trend in reducing senescence load induced by tobacco smoke in respiratory systems or renal ischemia-reperfusion in urinary systems (Ma *et al*, 2018; Wang *et al*, 2013) (**Table 3**).

### 3.5 Diet additives, single ingredient

#### 3.5.1 Nutrient

##### Amino acid

One study with low quality of methodology showed that L-citrulline and L-citrulline in combination with L-arginine had 1/2 positive and 1/2 insignificant outcome, while L-arginine had 2/2 insignificant outcomes in normal ageing (Tsuboi *et al*, 2018) (**Table 3**).

##### Carbohydrate

*Lycium barbarum* polysaccharides, investigated with moderate quality of methodology, demonstrated 2/6 positive outcomes with 2/6 outcomes showing a positive trend on three senescence markers in normal ageing (Xia *et al*., 2014). Alginate oligosaccharide had 3/3 positive outcomes reported with low quality of methodology in a D-galacotse-induced model (Feng *et al*, 2021) (**Table 3**).

##### Fat

In studies with moderate quality of methodology, the investigation of eicosapentaenoic acid (EPA) combined with docosahexaenoic acid (DHA) exhibited 7/13 positive outcomes (Qureshi *et al*, 2020), while corn oil in normal ageing showed 7/7 insignificant outcomes (Qureshi *et al*., 2020). Similarly, in a D-galactose-induced ageing model, arachidonic acid revealed 8/9 insignificant outcomes (Chen *et al*., 2017a) (**Table 3**).

##### Mineral

One study investigated the effect of phosphate binders lanthanum carbonate and calcium carbonate on four senescence markers, showing 3/6 and 2/6 positive outcomes respectively in a chronic kidney disease model (Yamada *et al*, 2015) (**Table 3**).

##### Vitamin

The effect of vitamin E and folic acid on senescence load was presented with high quality of methodology. Vitamin E had 15/21 positive outcomes reported in normal ageing (1 study) (Zeng *et al*., 2022) and D-galactose-induced ageing models (3 studies) (Chen *et al*., 2017a; Sun *et al*., 2020; Sun *et al*, 2018), while folic acid had 16/33 positive outcomes reported in normal ageing (1 study) (Ye *et al*, 2021b), sleep deprivation (1 study) (Zhang *et al*, 2019) and deficient folic acid models (2 studies) (Lv *et al*, 2019; Zhang *et al*, 2021c). The majority of outcomes displayed a positive trend (14/16) for vitamin C based on moderate quality of methodology (Jeong *et al*, 2018). Vitamin D showed 1/2 positive outcome and 1/2 negative outcome with low quality of methodology (Bima *et al*, 2021) (**Table 3**).

#### 3.5.2 Bioactives

##### Alkaloid

The investigation of caffeine with moderate quality of methodology yielded 9/13 positive outcomes in senescence models induced by ultraviolet irradiation (Li *et al*., 2018) and low density lipoprotein L5 (Wang *et al*, 2018). Matrine had 20/22 positive outcomes reported in a D-galactose-induced ageing model with moderate quality of methodology (Sun *et al*., 2018) (**Table 3**).

##### Glycoside

Studies with moderate quality of methodology showed that ginsenoside had 5/24 positive outcomes with 11/24 outcomes indicating a positive trend in normal ageing (Yu *et al*, 2020) and lead acetate-induced senescence models (Cai *et al*, 2018). Two positive outcomes out of four (2/4) in 2,3,5,40-tetrahydroxystilbene-2-O-b-D-glucoside (THSG) (Han *et al*, 2012) and 3/6 positive outcomes in salidroside (Xing *et al*, 2018) were reported with moderate quality of methodology (**Table 3**).

##### Lactone

The effect of yangonin on ethanol diet-induced senescence was investigated in two studies with moderate quality of methodology, revealing 27/27 positive outcomes (Dong *et al*, 2021; Kong *et al*, 2021) (**Table 3**).

##### Nucleoside and nucleotide

Nicotinamide riboside showed 9/18 positive outcomes in an Alzheimer’s disease transgenic mice model with moderate quality of methodology (Hou *et al*, 2021), while nicotinamide mononucleotide showed 4/4 positive outcomes in normal ageing with low quality of methodology (Huang *et al*, 2022) (**Table 3**).

##### Pigment

Studies with moderate quality of methodology demonstrated that lycopene had 6/6 positive outcomes in a D-galactose-induced ageing model (Liu *et al*, 2022); curcumol had 5/11 positive along with 6/11 outcomes showing a positive trend on senescence induced by high-fat diet (Qi *et al*., 2021); fucoxanthin had 4/6 insignificant outcomes on senescence in a sodium iodate-induced macular degeneration model (Chen *et al*, 2021) (**Table 3**).

##### Polyphenol

Three interventions were investigated with high quality of methodology on cellular senescence: resveratrol had 46/64 positive outcomes (8 studies) (Feng *et al*, 2020; Huang *et al*, 2017; Li *et al*, 2017; Liu *et al*, 2019; Liu *et al*., 2018; Liu *et al*., 2022; Luo *et al*, 2021; Xue *et al*, 2021); quercetin had 11/31 positive outcomes with 11/31 outcomes indicating a positive trend (3 studies) (Kim *et al*, 2019; Yu *et al*, 2021; Zhang *et al*., 2021a); fisetin had 26/39 outcomes with a positive trend (3 studies) (Alsuraih *et al*, 2021; Kim *et al*, 2021; Yousefzadeh *et al*, 2018). Studies with moderate quality of methodology reported positive outcomes in naringenin (8/8) (Hua *et al*, 2021), dihydromyricetin (12/12) (Qian *et al*, 2021) and puerarin (6/6) (Qian *et al*., 2021) as well as insignificant outcomes in coenzyme Q10 (14/18) (Tarry-Adkins *et al*, 2013) and epigallocatechin gallate (EGCG) (29/36) (Sharma *et al*, 2022) (**Table 3**).

##### Probiotic

One study with low quality of methodology showed an inconclusive effect of *Lactobacillus* strains on senescence induced by D-galactose with one or two positive outcomes out of four (Hor *et al*, 2019) (**Table 3**).

##### Pyrone

Maltol resulted in 12/12 positive outcomes in a D-galactose-induced ageing model with moderate quality of methodology (Sha *et al*, 2021) (**Table 3**).

##### Other

Studies with moderate quality of methodology showed that H2-rich water had 8/11 positive outcomes in mitigating senescence load induced by high-fat diet (Iketani *et al*, 2018) and cigarette smoke (Suzuki *et al*, 2017) (**Table 3**).

### 3.6 Human studies

Two interventions were investigated in four human studies. A high protein diet showed 1/2 insignificant outcome in p16 levels immediately post-exercise and 1/2 negative outcome in p16 levels 48 hours after exercise compared to a low protein diet with low quality of methodology (Yang *et al*, 2018). Three studies with high quality of methodology showed that glycoside ginsenoside Rg1decreased p16 but increased SASP (Interleukin-6) levels three hours after exercise (3/17 positive outcomes, 1/17 negative outcome) and showed no effect on SA-ß-gal and SASP (vascular endothelial growth factor) levels after exercise compared to placebo in healthy participants (13/17 insignificant outcomes) (Lee *et al*, 2021; Wu *et al*, 2020; Wu *et al*, 2019) (**Table 3**).

### 3.7 Interventions in the DrugAge database

Overall, 25 out of the total found 68 interventions had at least one entry in the DrugAge database (**Appendix B: Table S4**). The longevity extension property of resveratrol, quercetin, caffeine, vitamin E and EGCG had been investigated with at least ten entries.

### 3.8 Risk of bias

The SYRCLE’s risk of bias for animal studies is presented in **Fig 2A** and **Appendix B: Table S5**. Overall, 33 animal studies had a high risk of bias, whereas the rest 45 studies had an unclear risk of bias as none of the studies reported on allocation concealment, random housing, blinding of personnel and outcome assessor blinded. Two out of 78 studies had a low risk of bias on random outcome assessment. The main high risk of bias was derived from incomplete outcome data (26 studies) and selective outcome reporting reported (10 studies). Four out of 78 studies had a high risk in other sources of bias due to pooling samples without random selection (i.e. unit of analysis errors).

**Figure 2.**
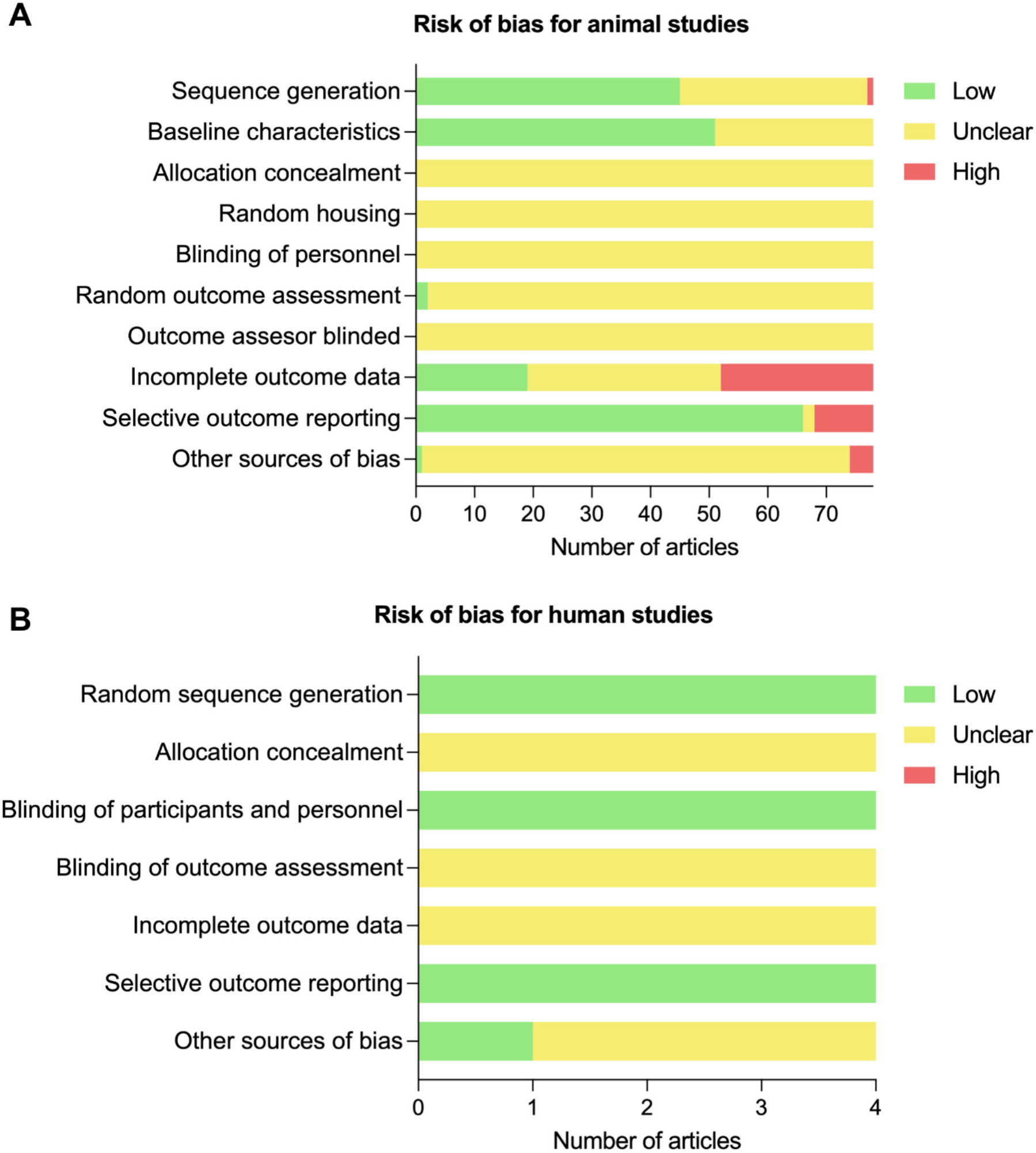
Risk of bias for (A) animal and (B) human studies using SYRCLE’s risk of bias tool and Cochrane risk of bias tool v2.0, respectively.

The Cochrane risk of bias for human studies is presented in **Fig 2B** and **Appendix B: Table S6**. All four studies had an overall unclear risk of bias, as none reported on allocation concealment, blinding of outcome assessment, and incomplete outcome data.

## Discussion

This comprehensive systematic review investigated the potential senotherapeutic association of diet and dietary ingredients in animal models and humans. Data obtained show that senescence can be targeted with senotherapeutics and single-ingredient diet additives resveratrol, vitamin E and soy protein isolate have the potential to reduce cellular senescence load in animal models. However, this finding should be interpreted with caution as eleven out of fifteen studies had an overall high risk of bias. Other interventions showed limited evidence as senotherapeutics in both animals and humans (**Fig 3**).

**Figure 3.**
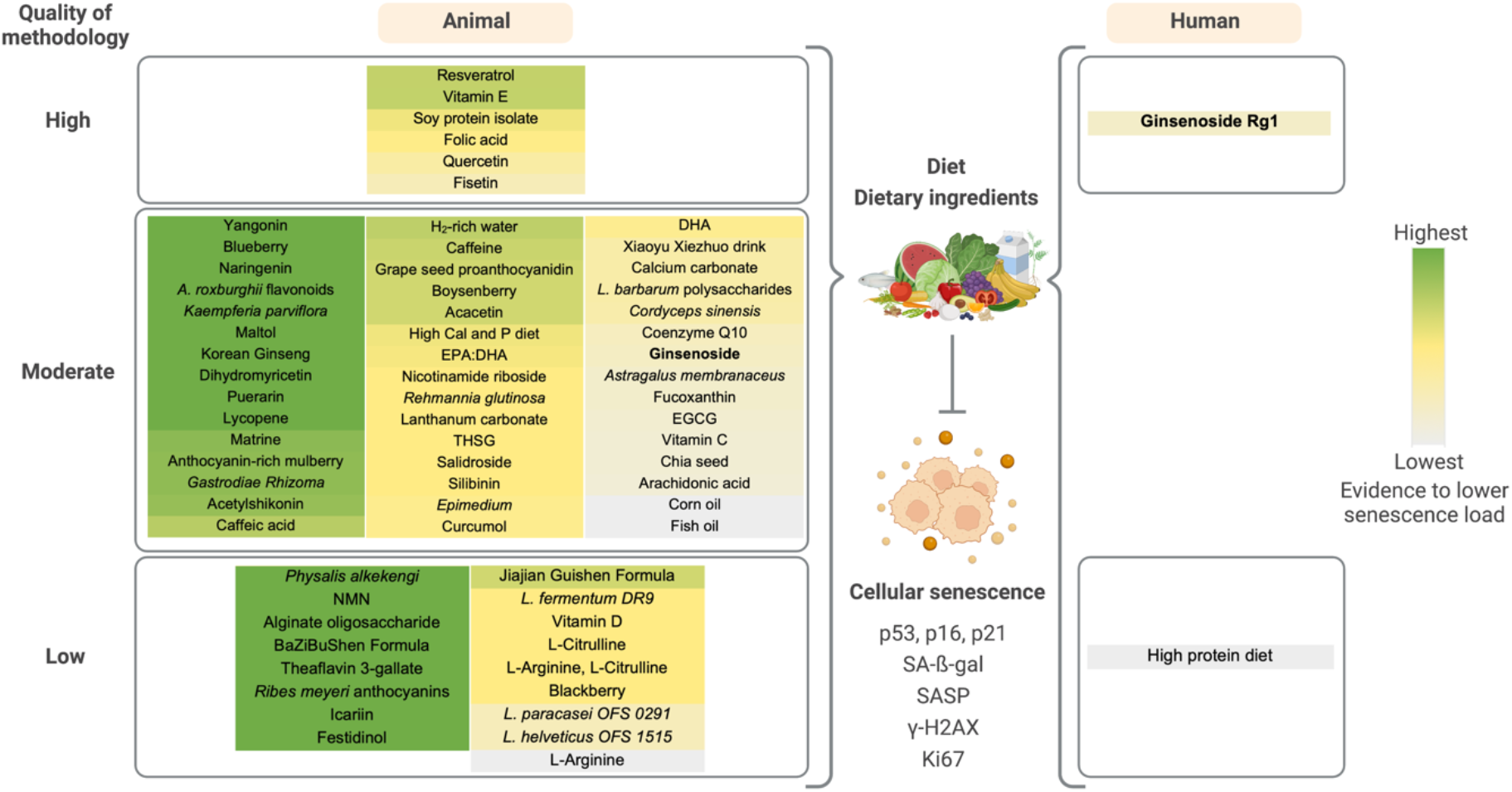
Overview of diet and dietary ingredients on reducing cellular senescence load in animals and humans stratified by the quality of methodology. Created with BioRender.com. Dietary interventions are ordered based on the evidence to lower senescence load. Quality of methodology: high, ≥ two senescence markers used in ≥ three studies; moderate, ≥ two senescence markers used in one or two studies; low, one senescence marker used. Abbreviations: *A. roxburghii* = *Anoectochilus roxburghii*, Cal = Calcium, DHA = Docosahexaenoic acid, EGCG = Epigallocatechin gallate, EPA = Eicosapentaenoic acid, *L.* = *Lactobacillus*., *L. barbarum* = *Lycium barbarum*, NMN = Nicotinamide mononucleotide, P = Phosphorus, SASP = Senescence-associated secretory phenotype, SA-ß-gal = Senescence-associated beta-galactosidase, THSG = 2,3,5,40-tetrahydroxystilbene-2-O-b-D-glucoside, γ- H2AX = H2A histone family member X phosphorylated on serine 139.

Resveratrol is a natural polyphenol inherent to berry fruits, such as grapes and blueberries, and peanuts among others. It is the most investigated dietary ingredient in either normal ageing (Liu *et al*., 2018), or senescence models induced by D-galactose (Li *et al*., 2017; Liu *et al*., 2022), ethanol liquid diet (Luo *et al*., 2021), high-fat diet (Xue *et al*., 2021), myocardial infarction surgery (Feng *et al*., 2020), overtraining (Liu *et al*., 2019), and concanavalin A (Huang *et al*., 2017). Orally taken resveratrol mitigates senescence load characterized by high levels of cell cycle regulators (p53, p21, p16, p19), SA-ß-gal, SASP and low levels of proliferation (Ki67, PCNA, Cyclin D2) in animal tissues across nervous, cardiovascular, digestive, and endocrine systems. This finding might partially be due to the pluripotent molecular effects of resveratrol which are cardioprotective, neuroprotective, anti-inflammatory, anti-hypertensive and anti-ageing (Zhang *et al*, 2021b). Vitamin E is a natural antioxidant primarily sourced from nuts, grains and olive oil (Boccardi *et al*, 2016). It was used as a positive control in four included studies in this current systematic review and was found to exert senotherapeutic effects on brain tissues from both normal ageing (Zeng *et al*., 2022) and D-galactose-induced senescence models (Chen *et al*., 2017a; Sun *et al*., 2020; Sun *et al*., 2018). This may suggest a protective role for vitamin E in brain ageing and neurogenerative diseases (La Fata *et al*, 2014). Soy protein is a vegetable protein isolated from soybean which exhibited benefits on musculoskeletal systems (Khairallah *et al*, 2017; Lin *et al*, 2018; Xiao, 2008). Three studies consistently showed that soy protein isolate had senotherapeutic effects on bone tissues from rat models with or without the presence of senescence inducers such as ovariectomy (Chen *et al*., 2017b; Zhang *et al*., 2014) and high-fat diet (Chen *et al*., 2015). Soy protein isolate is a composite of storage protein subunits, 7S and 11S (Nishinari *et al*, 2014). It is unclear which of the subunits is causing the observed effect.

It is crucial to acknowledge that the senotherapeutic effect of interventions is dependent on several factors. Firstly, the difference in genetic backgrounds of the mice used and the method of senescence induction might cause discrepancies. Oral intake of folic acid reduced senescence load in heart tissues from older mice but not in young mice (Ye *et al*., 2021b). Similarly, a high calcium and phosphate diet (Chen *et al*., 2019), epimedium (Zhao *et al*., 2022), vitamin C (Jiang *et al*, 2022), nicotinamide riboside (Hou *et al*., 2021), and fisetin (Alsuraih *et al*., 2021) were capable of reducing senescence load in transgenic mice models but not in wild-type mice. Vitamin D had opposite effects on senescence in rats fed on a high-carbohydrate diet and a normal diet (Bima *et al*., 2021). In a high-fat diet model that successfully induced senescence, quercetin was reported to display a senotherapeutic effect on kidney tissues (Kim *et al*., 2019), while in a high-fat diet model that failed to induce senescence, quercetin had no effect on SA-ß-gal, p21 and even increased the mRNA levels of p53, p16 and SASP in heart tissues (Yu *et al*., 2021). Moreover, the senotherapeutic effect varied in different biomaterials used. Vitamin E reduced senescence load induced by D-galactose in hippocampus, liver and spleen (Sun *et al*., 2020; Sun *et al*., 2018; Zeng *et al*., 2022) but not in testis (Chen *et al*., 2017a). In some cases, results were different even when the biomaterials originate from the same organ. Chia seed reduced senescence levels in epididymal adipose not in subcutaneous adipose (Rui *et al*., 2018); L-citrulline decreased SA-ß-gal activity in thoracic aorta not in abdominal aorta (Tsuboi *et al*., 2018); lactobacillus strains exerted different effects on musculoskeletal systems (e.g. *Lactobacillus fermentum DR9* had no effect on senescence in soleus muscle and positive effect in gastrocnemius muscle and tibia bone but showed a negative trend in femur bone) (Hor *et al*., 2019). Furthermore, an intervention can confer completely different effects at different dosages. Folic acid induced senescence at a dosage of 8 mg/kg diet in transgenic mice infected with hepatitis B (Zhang *et al*., 2021c). *Lycium barbarum* polysaccharides decreased SA-ß-gal levels in a dosage-dependent manner (dosage increased from 1 to 3 mg/mL), while it significantly increased SA-ß-gal activity at a dosage of 4 mg/mL compared to control in a zebrafish model (Xia *et al*., 2014).

Studies that were insufficiently reported or poorly designed, conducted and analysed had an unclear or high risk of bias, which pose a challenge to the translational reliability (Akhtar, 2015). For instance, fisetin showed a positive trend towards lowering senescence using p21, p16, ß-gal and SASP markers in transgenic mice, where the symbols for *p* values in the result figures were not explained in the legend (Yousefzadeh *et al*., 2018). Twenty-nine out of 78 studies were not primarily designed to investigate the senotherapeutic effect of interventions where senescence markers were measured as an underlying mechanism of alleviating poor conditions such as hematopoietic homeostasis defects (Cai *et al*., 2018), bone loss (Chen *et al*., 2017b), endothelial dysfunction (Furuuchi *et al*, 2018), Alzheimer’s disease (Hou *et al*., 2021), non-alcoholic steatohepatitis (Hua *et al*., 2021) or atherosclerosis (Serino *et al*, 2020). Studies that were not designed to investigate senescence as a primary aim often used one senescence marker. Normalization of age and sex at baseline is critical for senescence-related studies as senescence load is higher in tissues with higher chronological age (Tuttle *et al*., 2020) and different between female and male (Ng & Hazrati, 2022). Studies did not report age and/or sex, or only reported the age range of mice without comparing age between groups (Chen *et al*., 2021; Feng *et al*., 2020; Hong *et al*, 2021; Kim *et al*., 2021; Liu *et al*., 2019; Liu *et al*., 2022; Wang *et al*., 2013; Xing *et al*., 2018; Ye *et al*, 2021a; Zhang *et al*., 2021a; Zhang *et al*., 2019), resulting in an unclear risk of baseline characteristic bias. In 45 out of 78 studies, it was unclear if the allocation of mice was concealed, housing was random, and personnel who took care of animals and assessed outcomes was blinded. These are quintessential elements to consider during experimental evaluation and guarantee the quality of a study and the validity of results when conducting animal studies (Hooijmans *et al*., 2014).

More human studies that are better designed to address the potential senotherapeutic effect of diets or dietary ingredients are needed to facilitate the translation from animals. As there is no universal senescence marker, a combination of markers should be used. Lysosomal, proliferative features and senescence-associated genes concomitantly present with core SASP inflammatory markers have been proposed for validation of the presence and type of senescent state (Kohli *et al*, 2021). In addition, age and sex at baseline are recommended to be strictly controlled between groups to decrease inter-and intra-variability. Since obtaining human tissues is challenging, cellular senescence could be measured in human peripheral blood which is minimally invasive and holds sustainable advantages (Guan *et al*, 2022).

The present review synthesized current evidence about the effect of diet and dietary ingredients on senescence in animals and humans, providing evidence of promising interventions with potential senotherapeutic effects while the challenge of translation was highlighted. Some limitations need to be recognized. Firstly, the search terms specifically focused on ingredients in diets where caloric restriction was excluded as it is beyond the scope of this review. Secondly, SASP markers were not part of the search strategy as they are often measured in a non-senescent context. Thirdly, given that the heterogeneous type of senescence markers in tissues assessed by different techniques, a meta-analysis could not be performed to determine the overall senotherapeutic effect. Instead, the number of positive outcomes was counted based on reported *p* values which are largely influenced by sample size. Lastly, publication bias could not be ruled out as studies with positive results were likely to be published whereas negative results were not. Despite these limitations, this is the most comprehensive systematic review that provides solid evidence for the association of the dietary ingredients identified in animal models as senotherapeutics.

## Conclusion

Resveratrol, vitamin E and soy protein isolate are promising senotherapeutics in animal models and should be translated into humans.

## Acknowledgements

The authors thank the librarian Annelissa Mien Chew Chin from National University of Singapore Medical Library for her assistance in the search strategy.

## Appendix A

Table S1. Inclusion and exclusion criteria.

Search Strategy

## Appendix B

Table S1. Cellular senescence markers eligible for data extraction with expected change in direction of outcome after an intervention.

Table S2. Classification of interventions aiming to reduce cellular senescence in animals and humans.

Table S3. Classification of biomaterials used in studies into physiological organ systems.

Table S4. Interventions with at least one entry in DrugAge database.

Table S5. SYRCLE’s risk of bias for animal studies, stratified by the classification of interventions.

Table S6. Cochrane risk of bias for human studies.

## Appendix C

Table S1. Characteristics of intervention studies in animals, stratified by the classification of interventions.

Table S2. Methods and results of intervention studies in animals, stratified by the classification of interventions.

Table S3. Characteristics of intervention studies in humans, stratified by the classification of interventions.

Table S4. Methods and results of intervention studies in humans, stratified by the classification of interventions.

